# Lexical influences on categorical speech perception are driven by a temporoparietal circuit

**DOI:** 10.1101/2020.08.11.246793

**Authors:** Gavin M. Bidelman, Claire Pearson, Ashleigh Harrison

## Abstract

Categorical judgments of otherwise identical phonemes are biased toward hearing words (i.e., “Ganong effect”) suggesting lexical context influences perception of even basic speech primitives. Lexical biasing could manifest via late stage post-perceptual mechanisms related to decision or alternatively, top-down linguistic inference which acts on early perceptual coding. Here, we exploited the temporal sensitivity of EEG to resolve the spatiotemporal dynamics of these context-related influences on speech categorization. Listeners rapidly classified sounds from a /gi/ - /ki/ gradient presented in opposing word-nonword contexts (*GIFT-kift* vs. *giss-KISS*), designed to bias perception toward lexical items. Phonetic perception shifted toward the direction of words, establishing a robust Ganong effect behaviorally. ERPs revealed a neural analog of lexical biasing emerging within ∼200 ms. Source analyses uncovered a distributed neural network supporting the Ganong including middle temporal gyrus (MTG), inferior parietal lobe (IPL), and middle frontal cortex. Yet, among Ganong-sensitive regions, only left MTG and IPL predicted behavioral susceptibility to lexical influence. Our findings confirm lexical status rapidly constrains sub-lexical categorical representations for speech within several hundred milliseconds but likely does so outside the purview of canonical “auditory-linguistic” brain areas.

## 1. Introduction

An important building block for language is the ability to transform sensory information into abstract linguistic representations (Goldstone and Hendrickson, 2010). Speech sounds vary continuously across time, environments, speaker identities, and stimulus contexts, and yet, listeners easily parse the speech stream into discrete phonemes (Lotto and Holt, 2016; Phillips, 2001; Pisoni and Luce, 1987). The categorical perception (CP) of speech maps infinitely variable acoustic signals into discrete phonetic-linguistic representations on which the speech-language system can operate (Liberman et al., 1967; Pisoni, 1973; Pisoni and Luce, 1987). CP is indicated when gradually morphed speech sounds along a continuum are heard as belonging to one of a few discrete phonetic classes. Tokens labeled with different identities are said to cross the categorical boundary, a psychological border where listeners’ responses abruptly flips due to a perceptual warping of the stimulus space (i.e., compression of within-category sounds) (Best and Goldstone, 2019; Goldstone et al., 2000; Livingston et al., 1998).

One nebulous issue in speech perception concerns whether higher-level activation of lexical representations directly affects sub-lexical components (e.g., phoneme categories). On one extreme is the ridged view that once established, internalized speech prototypes (i.e., equivalence classes or category members) are invariant to superficial stimulus manipulation or lexical context (Liberman et al., 1957). Under this model, categories are impervious to influences from surrounding information and sound elements that precede or follow an isolated stimulus cannot influence its categorization or location of the perceptual boundary. On the contrary, acoustic-phonetic categories—traditionally considered early or lower-level constructs of the speech signal—are in fact highly malleable to contextual variations (Elman and McClelland, 1988; Francis and Ciocca, 2003; Ganong, 1980; Holt and Lotto, 2010; Myers and Blumstein, 2008; Norris et al., 2003; Pisoni, 1975). Moreover, the degree to which context influences the category identity of speech varies with language experience (Bidelman and Lee, 2015; Kuhl et al., 1992; Lively et al., 1993). Consequently, it is now well-established that phonetic categories are flexible and perception of even individual speech features depends critically on the surrounding signal (Repp and Liberman, 1987).

Context-dependent effects in CP are best illustrated by the so-called “Ganong effect” (Ganong, 1980). The Ganong phenomenon occurs when listeners’ perceived category boundary of a word-nonword continuum of phonemes shifts (is biased) towards the lexical item. When perceiving a “da-ta” continuum, for example, English-speaking listeners show a stark shift in their perceptual category boundary towards lexical items when one of the gradient’s endpoints contains a real word (e.g., “DASH-tash”) (Ganong, 1980; Ganong and Zatorre, 1980). Similar interpretive biasing can be induced via learning when listeners are exposed to new contexts that shapes their perception of otherwise isolated sounds (Norris et al., 2003). Collectively, behavioral studies suggest that stimulus context expands the mental category for expected or behaviorally relevant stimuli (McMurray et al., 2008).

One interpretation of lexical effects is that they reflect direct linguistic influence on perceptual processes. Alternatively, another school of thought argues lexical context effects are post-perceptual and are therefore related to executive mechanisms (i.e., response selection, decision). Fox (1984) tested the interaction between lexical knowledge and phonetic categorization during speech perception using Ganong-like stimuli. Lexical status did not influence phonetic categorization at shorter response latencies or when participants were given a response deadline, suggesting lexical context influences later stimulus selection rather than perceptual encoding, *per se*. This notion is supported by results from Pitt and Samuel (1993), who found the strength of lexical influences on perception of ambiguous sound tokens depended on their position in a word—lexical effects were weaker when tokens occurred toward the beginning compared to the end of words. These data support “late stage” or “selection-based” models whereby the very formation of categories themselves only emerge at the late decision-stage of the processing hierarchy (e.g., MERGE model; Norris et al., 2000).

Rather than acting at late stages, lexical biasing could instead manifest via top-down (and perhaps bi-directional) modulations of early perceptual processing with the lexical interface. Indeed, growing evidence from neuroimaging studies (Gow et al., 2008; Myers and Blumstein, 2008; Noe and Fischer-Baum, 2020; van Linden et al., 2007) reaffirms such interactive, connectionist views of categorization (e.g., TRACE; McClelland and Elman, 1986). Employing fMRI with a Ganong task, Myers and Blumstein (2008) found that the placement of the phonetic boundary modulated activity both in perceptual [e.g., superior temporal gyrus, STG), inferior parietal lobule (IPL)] and frontal executive brain areas (IFG, ACC), with greater activity for ambiguous items near the boundary. The mere involvement of the STG strongly suggested lexical shifts not due solely to executive decision processes but at minimum, includes a perceptual component that either itself has direct access to lexical properties or is interactively reactivated to integrate phonetic and extra-phonetic factors in placing the phonetic boundary (Gow et al., 2008; Myers and Blumstein, 2008; Noe and Fischer-Baum, 2020). While fMRI offers excellent spatial characterization of potential lexical effects, it lacks the temporal precision necessary to resolve the underlying brain dynamics of category formation (Bidelman et al., 2013) and related lexical influences (Gow et al., 2008), both of which unfold within a few hundred milliseconds after speech onset (e.g., Mahmud et al., 2020).

Extending prior neuroimaging work (Gow et al., 2008; Myers and Blumstein, 2008), the aim of the present study was to characterize the spatiotemporal dynamics of context-dependent lexical influences on CP with the goal of establishing where and when speech categories are prone to Ganong-like biasing. We used EEG coupled with source reconstruction to assess the underlying neural bases of phoneme categorization and its lexical modulation. Our task included word-nonword (*GIFT-kift*) and nonword-to-word (*giss-KISS*) acoustic gradients of an otherwise identical /gi/-/ki/ acoustic-phonetic continuum designed to bias listeners’ perception toward the lexical item and shift their perceptual category boundary (Ganong, 1980; Myers and Blumstein, 2008). Our findings confirm that lexical status rapidly (∼200-300 ms) constrains sub-lexical category speech representations but further suggests this interactivity occurs outside canonical auditory-linguistic brain structures. Instead, among Ganong-sensitive brain regions, we find engagement of a temporoparietal circuit (i.e., inferior parietal, middle temporal gyrus) is critical to describing listeners’ susceptibility to contextual biasing during category judgements.

## 2. Materials & Methods

### 2.1 Participants

Sixteen young adults (3 male, 13 female; age: *M* = 24.5, *SD* = 12.9 years) were recruited from the University of Memphis student body^1^. Sample size was based on several previous neuroimaging studies on context-effects in CP (e.g., Gow et al., 2008; Myers and Blumstein, 2008). All exhibited normal hearing sensitivity confirmed via audiometric screening (i.e., < 25 dB HL, octave frequencies 250 - 8000 Hz). Each participant was strongly right-handed (74.8 ± 27.0% laterality index; Oldfield, 1971), had obtained a collegiate level of education (18.8 ± 2.7 years formal schooling), and was a native speaker of American English. Participants were considered nonmusicians (e.g., Mankel and Bidelman, 2018), having on average 3.25 ± 3.3 years of music training. All were paid for their time and gave informed consent in compliance with a protocol approved by the Institutional Review Board at the University of Memphis.

### 2.2 Speech stimulus continua

Stimuli were adapted from Myers and Blumstein (2008). Speech tokens consisted of a /gi/ to /ki/ stop-consonant continuum presented in two word/nonword contexts. Each continuum was constructed using 8 equally spaced voice-onset times (VOTs) incrementing from 18 ms (/g/ percept) to 70 ms (/k/ percept) (**Fig. 1**). This otherwise identical VOT continuum was used to create word-to-nonword (*GIFT-kift*) and nonword-to-word (*giss-KISS*) gradients designed to bias listeners’ phonemic perception toward the lexical item (**Fig. 1b**). This was achieved by splicing the appropriate aspiration (i.e., “-ft” for *GIFT-kift;* “-ss” for *giss-KISS*) to the end of the otherwise identical /gi/-/ki/ sounds (for details, see Myers and Blumstein, 2008). All tokens were 200 ms in duration and RMS amplitude normalized.

**Figure 1:**
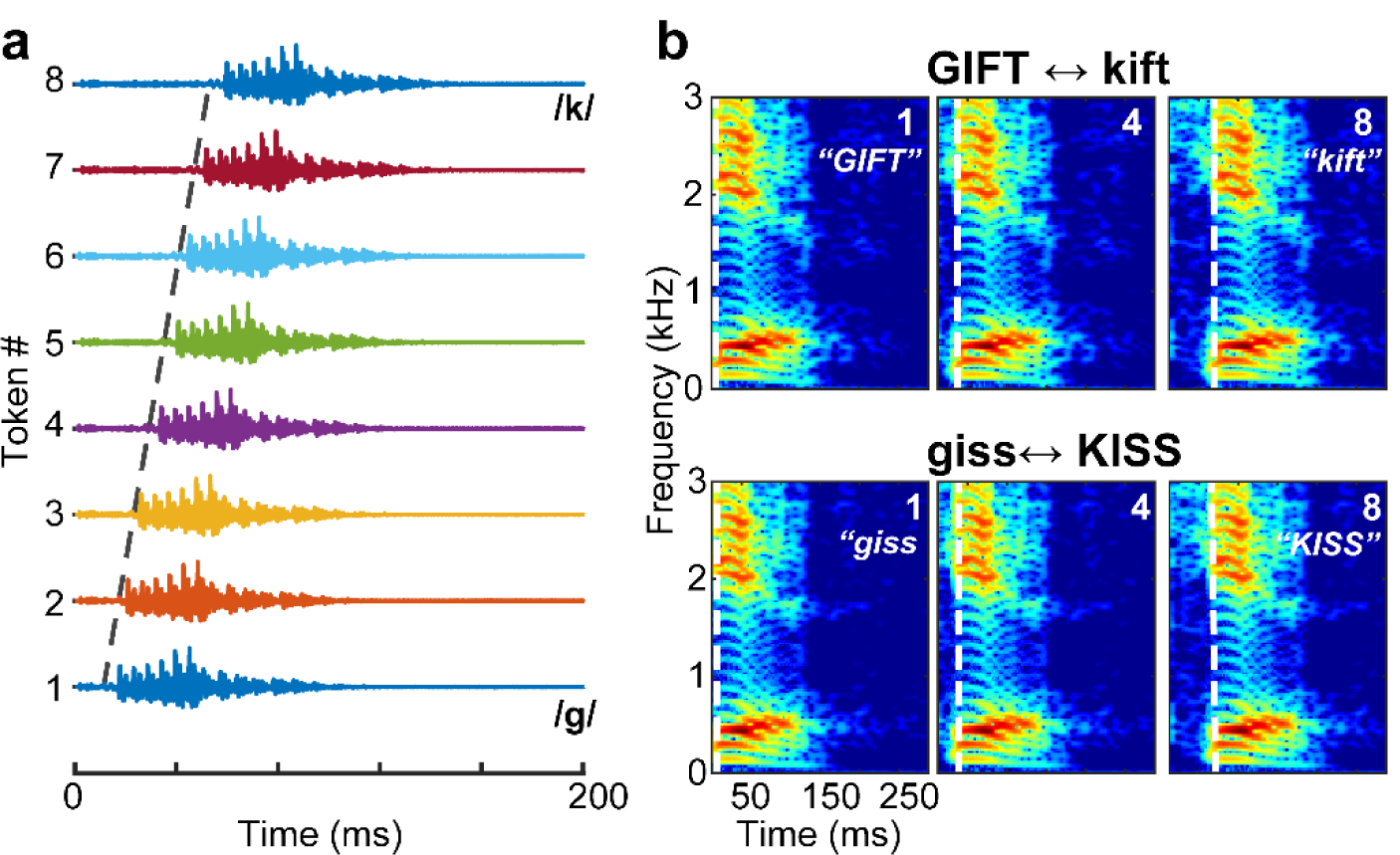
Speech stimuli used to probe the neural basis of lexical effects on categorical speech processing. (**a**) Acoustic waveforms of the continuum. Stimuli varied continuously in equidistant VOT steps to yield a morphed gradient from /gi/ to /ki/. (**b**) Spectrograms. The /gi/ to /ki/ continuum was presented in one of two word-nonword contexts (*GIFT-kift* and *giss-KISS*) such that at any point along the acoustic gradient, the same stop consonant could be perceived more as a word (or nonword) depending on lexical bias from the continuum’s endpoint. Dotted lines, onset of voicing demarcating VOT duration.

During EEG recording, listeners heard 120 trials of each individual token (per context) in which they labelled the sound with a binary response (“g” or “k”) as quickly and accurately as possible. Following, the interstimulus interval (ISI) was jittered randomly between 800 and 1000 ms (20 ms steps, uniform distribution) to avoid rhythmic entrainment of the EEG and anticipating subsequent stimuli. Block order for the *GIFT-kift* vs. *giss-KISS* continua were randomized within and between participants. The auditory stimuli were delivered binaurally at 79 dB SPL through shielded insert earphones (ER-2; Etymotic Research) controlled by a TDT RP2 signal processor (Tucker Davis Technologies).

### 2.3 EEG recordings

EEGs were recorded from 64 sintered Ag/AgCl electrodes at standard 10-10 scalp locations (Oostenveld and Praamstra, 2001). Continuous data were digitized at 500 Hz (SynAmps RT amplifiers; Compumedics Neuroscan) using an online passband of DC-200 Hz. Electrodes placed on the outer canthi of the eyes and the superior and inferior orbit monitored ocular movements. Contact impedances were maintained < 10 kΩ. During acquisition, electrodes were referenced to an additional sensor placed ∼ 1 cm posterior to Cz. Data were re-referenced offline to the common average for analysis. Pre-processing was performed in BESA® Research (v7.1) (BESA, GmbH). Ocular artifacts (saccades and blinks) were corrected in the continuous EEG using principal component analysis (PCA) (Picton et al., 2000). Cleaned EEGs were then filtered (1-20 Hz), epoched (−200-800 ms), baselined to the pre-stimulus interval, and ensemble averaged resulting in 16 ERP waveforms per participant (8 tokens*2 contexts).

### 2.4 Behavioral data analysis

Identification scores were fit with a sigmoid function *P* = 1/[1+*e*^-*β1*(*x* - *β0*)^], where *P* is the proportion of trials identified as a given vowel, *x* is the step number along the stimulus continuum, and *β*_*0*_ and *β*_*1*_ the location and slope of the logistic fit estimated using nonlinear least-squares regression.Comparing parameters between speech contexts revealed possible differences in the “steepness” (i.e., rate of change) and more critically, the location of the categorical boundary as a function of speech context. A lexical bias (i.e., Ganong effect) is indicated when the location of the perceptual boundary (*β*_*0*_) in phoneme identification shifts dependent on the anchoring speech context (Ganong, 1980; Myers and Blumstein, 2008). Behavioral labeling speeds (i.e., reaction times [RTs]) were computed as listeners’ median response latency across trials for a given condition. RTs outside 250-2500 ms were deemed outliers (e.g., fast guesses, lapses of attention) and were excluded from the analysis (Bidelman et al., 2013; Bidelman and Walker, 2017).

### 2.5 EEG data analysis

#### ERP sensor responses

From channel-level waveforms, we measured lexical bias effects in the speech ERPs by comparing scalp topographies at the ambiguous midpoint token (Tk4) evoked in the two different speech contexts (i.e., GIFT4 vs. KISS4). This token step is where lexical bias effects were most prominent behaviorally (see Fig. 2). Topographic t-tests were conducted in EEGLAB (Delorme and Makeig, 2004).

**Figure 2:**
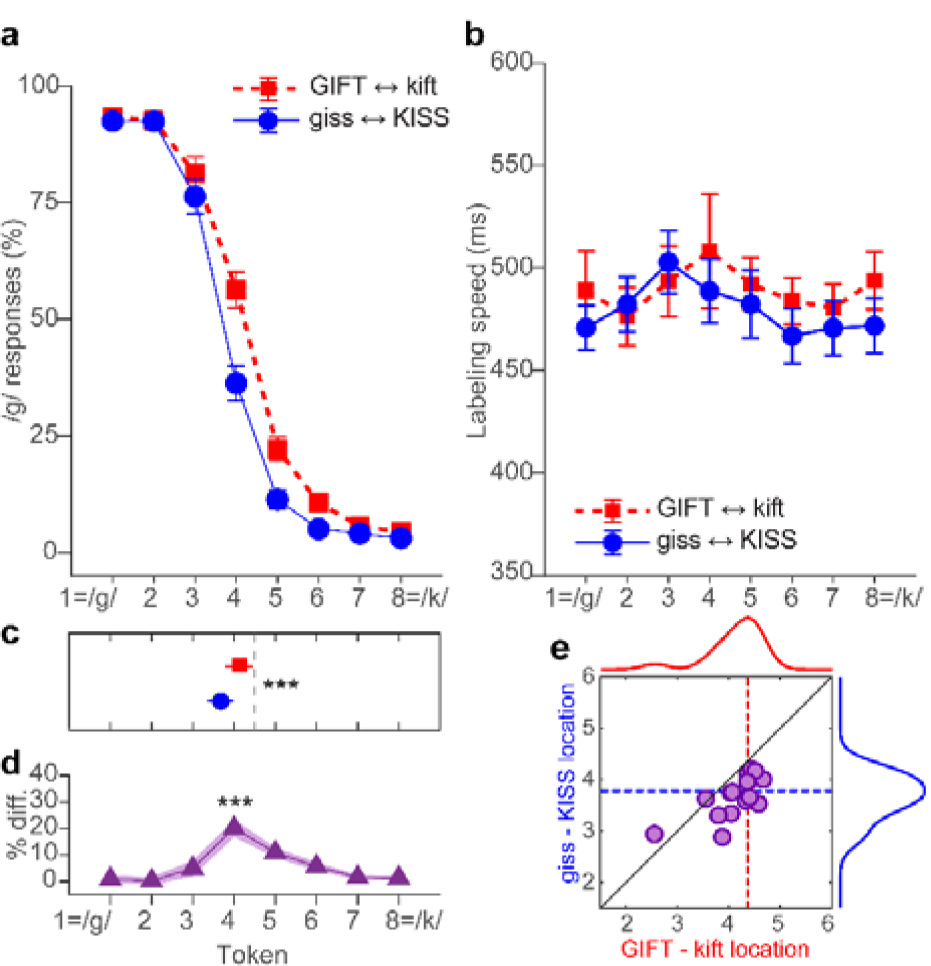
Lexical context biases the perceptual categorization of speech. (**a**) Psychometric identification functions show a shift in the perceptual boundary towards lexical items. Listeners more frequently reported /g/ responses in the *GIFT-kift* continuum and more /k/ responses for the *giss-KISS* context, confirming perception for otherwise identical stop consonants is biased toward hearing words. (**b**) Reaction times. Labeling speeds are faster for endpoint vs. midpoint tokens of the continuum consistent with category ambiguity near the midpoint of the continuum (Pisoni and Tash, 1974). (**c**) Critically, the location of the perceptual boundary (i.e., *β*_0_) shifts depending on the lexical context. (**d**) Identification performance differs maximally between contexts near the midpoint of the continua (i.e., Tk 4). (**e**) Comparison of boundary locations (*β*_0_) for the *GIFT-kift* vs. *giss-KISS* continua. The diagonal represents the case of an identical perceptual boundary between contexts. Boundaries shift leftward for *giss-KISS* compared to *GIFT-kift*, reflecting a higher precedence of /k/ responses in that context (vice versa for the other context). errorbars = ±1 s.e.m; *\*\*\*p<*0.0001.

#### Source analysis

To estimate the underlying sources contributing to the lexical effect, we used Classical Low Resolution Electromagnetic Tomography Analysis Recursively Applied (CLARA) [BESA® (v7)] (Iordanov et al., 2014) to estimate the neuronal current density underlying the scalp ERPs (e.g., Alain et al., 2017; Bidelman, 2018). CLARA models the inverse solution as a large collection of elementary dipoles distributed over nodes on a mesh of the cortical volume. The algorithm estimates the total variance of the scalp data and applies a smoothness constraint to ensure current changes minimally between adjacent brain regions (Michel et al., 2004; Picton et al., 1999). CLARA renders more focal source images by iteratively reducing the source space during repeated estimations. On each iteration (x2), a spatially smoothed LORETA solution (Pascual-Marqui et al., 2002) was recomputed and voxels below a 1% max amplitude threshold were removed. This provided a spatial weighting term for each voxel on the subsequent step. Two iterations were used with a voxel size of 7 mm in Talairach space and regularization (parameter accounting for noise) set at 0.01% singular value decomposition. Source activations were visualized on BESA’s adult brain template (Richards et al., 2016).

#### Dipole source modeling

To quantify the time course of source activations, we seeded discrete dipoles within the activation centroids identified in the CLARA volume images at a latency of 286 ms, where scalp data showed maximally lexical effects (see Fig. 4a) (cf. Bidelman and Walker, 2019). CLARA localized activity to five major foci including middle temporal gyrus (MTG), inferior parietal lobe (IPL), and middle frontal gyrus (MFG) in left hemisphere, and precentral gyrus (PCG) and insular (INS) cortex of right hemisphere (see Fig. 4d). Dipole time courses represent the estimated current within each regional source. We then used this 5 dipole model to create a virtual source montage to transform each participant’s scalp potentials (sensor-level recordings) into source space (Scherg et al., 2019; Scherg et al., 2002). This digital re-montaging applied a spatial filter to all electrodes (defined by the foci of our dipole configuration) to transform the electrode recordings to a reduced set of source signals reflecting the neuronal current (in units nAm) as seen *within* each anatomical region of interest (ROI) (Bidelman, 2018; Bidelman et al., 2018). Critically, we fit individual dipole orientations to each participant’s own data (anatomical locations remained fixed) to maximize the explained variance of the model at the individual subject level. The model provided a good fit to the grand averaged scalp data (Goodness of fit, entire epoch window = 75%), confirming the ERPs could be described by a restricted number of sources.

#### Brain-behavior correspondence

From the source waveform time courses, we measured peak amplitudes within the 200-300 ms time window, where lexical effects were prominent in raw EEG data (see Fig 4a,b). We then regressed source amplitudes (for each ROI) with listeners’ behavioral Ganong effect, computed as the magnitude of shift in their perceptual boundary between speech contexts (i.e., data in Fig. 2c). The allowed us to assess the behavioral relevance of each brain ROI and how context-dependent changes in neural activity (i.e., “neural Ganong” effect) relate to lexical biases in CP measured behaviorally.

### 2.6 Statistics

We analyzed the data using mixed-model ANOVAs in R (R Core team, 2018; *lmer4* package) with fixed effects of token (8 levels) and speech context (2 levels). Subjects served as a random effect. Multiple comparisons were corrected using Tukey-Kramer adjustments. Brain-behavior relations were assessed using robust regression (bisquare weighting) performed using the ‘fitlm’ function in MATLAB® 2020a (The MathWorks, Inc.).

## 3. Results

### 3.1 Behavioral data

Behavioral identification functions are shown for the two speech contexts in **Figure 2a**. Listeners more frequently reported /g/ responses in the *GIFT-kift* continuum and more /k/ responses for the *giss-KISS* context, confirming that perception for otherwise identical stop consonants is biased toward hearing words. The perceptual boundary location depended strongly on context [*t*_*15*_ *=* 2.37, *p* < 0.0317] (**Fig. 2c** and **2e**). Consistent with prior studies (Ganong, 1980; Myers and Blumstein, 2008; Noe and Fischer-Baum, 2020), context-dependent effects in CP where most evident near the ambiguous midpoint of the continuum (Tk 4), where listeners identification abruptly shifted phoneme categories [*t*_*15*_ *=* 6.00, *p* < 0.0001] (**Fig. 2d**). Ganong shifts also varied across individuals (e.g., Lam et al., 2017), with some listeners showing strong influence to lexical bias and others showing little to no changes in perception with speech context (**Fig. 3**).

**Figure 3:**
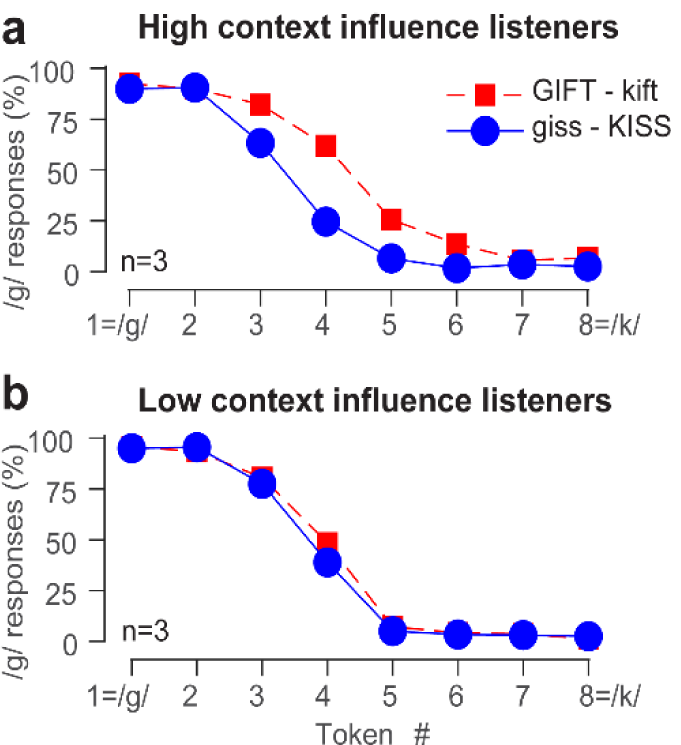
Lexical influences on CP are subject to individual differences. Identification functions for representative listeners (n=3) who showed the strongest (a) and weakest (b) influence of lexical context on speech categorization. High influence listeners perceptual boundary shifts dramatically with context whereas low influence listeners show little change in perception with lexical context.

**Figure 4:**
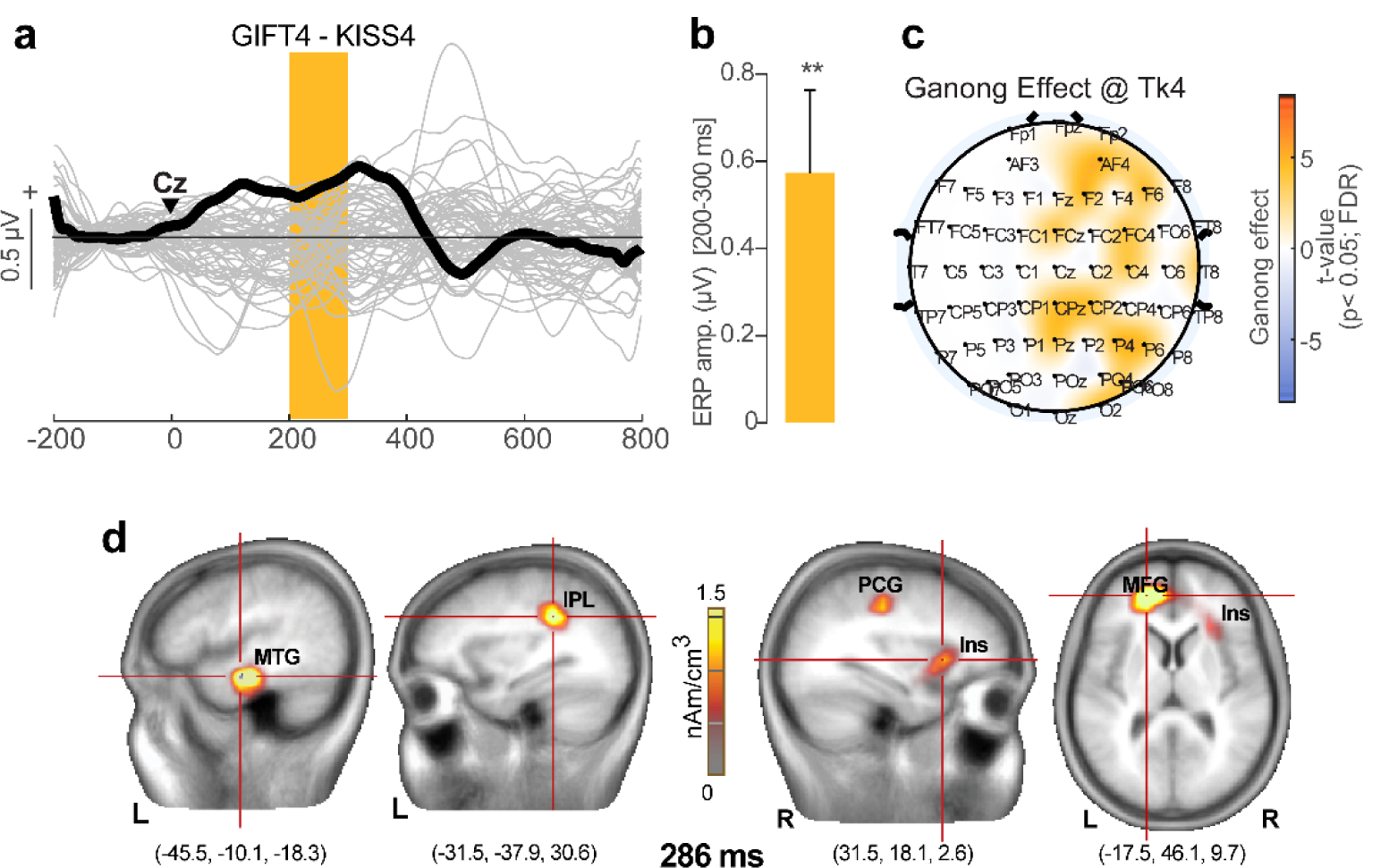
Neuroelectric brain activity reveals evidence of lexical biasing on speech categories. **(a)** Butterfly plot of ERP time courses (bold = Cz electrode) reflecting difference waves between Tk 4 responses when presented in *GIFT-kift* vs. *giss-KISS* contexts. ▾= speech stimulus onset. A running t-test (Guthrie and Buchwald, 1991) reveals lexical biasing between 200-300 ms (*p*<0.05; shaded segment). **(b)** Mean ERP amplitude at Cz (200-300 ms) differs from 0 indicating differentiation of identical speech tokens dependent on lexical context. (**c**) Topographic distribution of the Ganong effect across the scalp. Statistical maps (paired t-test, p<0.01, FDR-corrected; Benjamini and Hochberg, 1995) contrasting Tk4 responses in the two contexts. (**d**) Brain volumes show CLARA (Iordanov et al., 2014) distributed source activation maps underlying lexical bias during speech catherization. Maps were rendered at latency of 286 ms, where the effect was most prominent at the scalp (e.g., Fig 4a). Functional data are overlaid on an adult template brain (Richards et al., 2016). MTG, middle temporal gyrus, IPL, inferior parietal lobe, (MFG) middle frontal gyrus, (PCG) precentral gyrus, (INS) insular cortex. errorbars = ±1 s.e.m.; \*\**p*<0.01.

Speech labeling speeds were strongly modulated by context [*F*_*1,225*_ = 5.15, *p*=0.024] and token [*F*_*7,225*_ = 2.14, *p*=0.0408] (**Fig. 2b**). Identification was faster overall when categorizing tokens in the *giss-KISS* context (*p*=0.024). *GIFT-kift* responses further showed the hallmark slowing near the midpoint of the continuum relative to prototypical endpoints [Tk 1,2,7,8 vs. Tk 4,5: *p* =0.024] (Bidelman and Walker, 2017; Pisoni and Tash, 1974), attributable to more ambiguity in decision nearer the perpetual boundary. This categorical RT pattern was not observed for *giss-KISS* (*p* = 0.14). Collectively, these behavioral results suggest that lexical information (words) biases listeners’ categorization of otherwise identical phonetic features; even basic phoneme perception is latticed by the surrounding lexical context of the speech signal.

**Figure 5:**
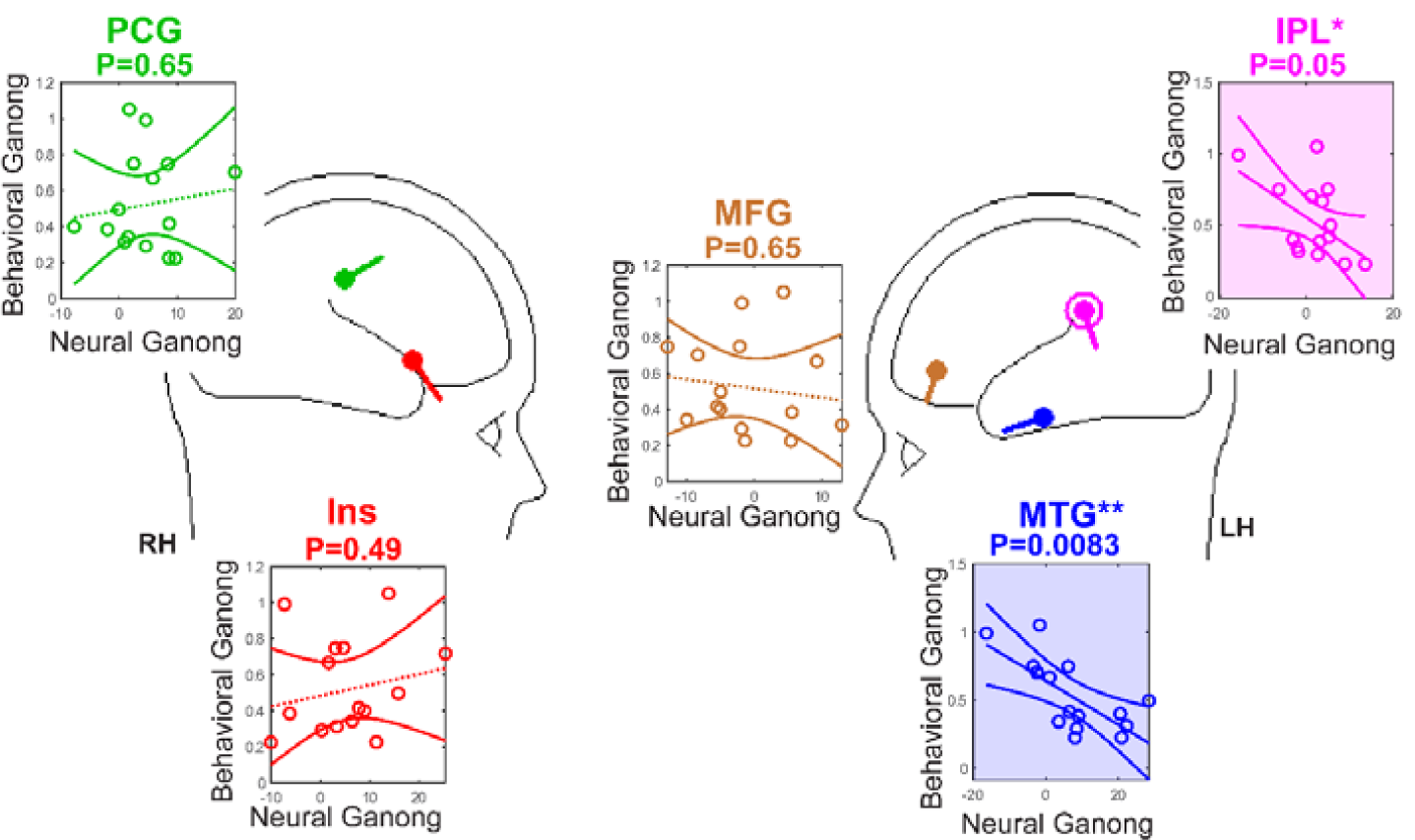
Lexical bias in CP is driven by engagement of MTG and IPL in left hemisphere. Cartoon heads illustrate the location of the dipole sources underlying the neural Ganong effect. Individual scatters show the relation between neural and behavioral Ganong effect measured from each ROI (shading, *p* <0.05). Solid regression lines, significant brain-behavior relation; Dotted lines, *n*.*s*. Flanking curved lines reflect 95% CIs. Of the active regions, only left MTG and IPL correspond with listeners’ behavioral bias. MTG, middle temporal gyrus, IPL, inferior parietal lobe, (MFG) middle frontal gyrus, (PCG) precentral gyrus, (INS) insular cortex, LH/RH, left/right hemisphere. \**p*<0.05; \*\**p*<0.01.

### 3.2 Electrophysiological data

Scalp ERPs are shown in **Figure 4**. To quantify the “neural Ganong” effect we contrasted ERPs to tokens at the perceptual boundary (i.e., Tk 4) (e.g., Myers and Blumstein, 2008), where lexical bias was strongest behaviorally (see Fig 2). Difference waves computed between midpoint tokens evoked during *giss-KISS* vs. *GIFT-kift* continua revealed context-dependent modulations in the time window between 200-300 ms [*t*_14_ = 3.03, *p*=0.009] (**Fig. 4a, b**). That is, despite identical acoustic information, stop consonants were perceived differentially depending on the word context they carried. The topography of the neural Ganong was broadly distributed over the scalp, spanning frontal, temporal, and parietal electrodes (**Fig. 4c**).

Source analysis of the ERPs exposed neural activations coding lexical bias in CP within five major foci among the auditory-linguistic-motor loop (e.g., Hickok and Poeppel, 2007; Rauschecker and Scott, 2009), including MTG, IPL (proximal to supramarginal gyrus, SMG), and MFG in left hemisphere, and PCG and INS in right hemisphere (**Fig. 4d**). For each participant, we extracted the time course of source activity from dipoles seeded at the centroids of these ROIs. We then measured and regressed the peak activation within each ROI—reflecting the magnitude of “neural Ganong”—against listeners’ behavioral Ganong (i.e., perceptual boundary shift; Fig. 2c). These brain-behavior correlations revealed strong associations between left MTG and left IPL activity and behavioral bias. These findings suggest that context-dependent modulations within a restricted temporo-parietal circuit were most inducive to describing the degree to which listeners’ CP was susceptible to lexical influences.

## Discussion

By measuring neuroelectric brain activity during rapid speech categorization tasks, our data reveal strong lexical bias in phonetic processing; perception for otherwise identical speech phonemes is attracted toward the direction of words, shifting listeners’ categorical boundary dependent on surrounding speech context. We show a neural analog of lexical biasing emerging within ∼200 ms within brain activity localized to a distributed, bilateral temporoparietal network including MTG and IPL. Our findings confirm that when perceiving speech, lexical status rapidly constrains sub-lexical representations to their category membership within several hundred milliseconds, establishing a direct linguistic influence on early speech processing.

Decoding speech and lexical biasing could be realized via phonetic “feature detectors” (Eimas and Corbit, 1973) that occupy and are differentially sensitive to different segments of the acoustic-linguistic space. Tunable detectors would tend to create quasi “acoustic foveae” that naturally build categories via overrepresentation of the stimulus space near stimulus protypes (Rozsypal et al., 1985). Adaptation studies—in which continuum sounds are presented repetitively and/or in serial order (Eimas and Corbit, 1973; Miller, 1975)—suggest that movement of the category boundary is explained by one detector becoming more desensitized from fatigue, thereby causing a boundary shift in the direction toward the un-adapted detector at the polar end of the continuum (Rozsypal et al., 1985). As confirmed empirically, larger boundary shifts would be expected for less strongly categorized continua (Rozsypal et al., 1985), e.g., vowels vs. stop consonants (Altmann et al., 2014), acoustically degraded speech (Bidelman et al., 2020; Bidelman et al., 2019), and for ambiguous speech tokens as shown here and previously (Ganong, 1980; Gow et al., 2008; Lam et al., 2017; Myers and Blumstein, 2008; Noe and Fischer-Baum, 2020). Alternatively, the Ganong-like displacements in perception we observe could occur if linguistic status moves the category boundary toward the most likely lexical candidate. Similarly, nonlinear dynamical models of perception posit that lexical items more strongly activate perceptual “attractor states” which pull auditory percepts toward word items (Tuller et al., 1994). Under this interpretation, the brain would presumably process even non-speech sounds through the lens of a “speech mode,” continually monitoring the auditory stream for lexical units (Liberman et al., 1981; Remez et al., 1981).

Considerable debate persists as to whether lexical effects in spoken word recognition result from feedback or feedforward processes (Gow et al., 2008; Myers and Blumstein, 2008; Norris et al., 2000; Pitt, 1995; Samuel and Pitt, 2003). Ganong shifts could occur if lexical knowledge exerts top-down influences to directly affect perceptual processes. Under these frameworks, lexical-based modulation of auditory-sensory brain areas (i.e., STG; Myers and Blumstein, 2008; van Linden et al., 2007) could result from top-input from higher levels associated with wordforms (e.g., SMG, MTG). Alternatively, a purely feedforward architecture (Norris et al., 2000) posits that lexical and phonetic outputs combine and interact at later post-perceptual stages of processing that are intrinsic to overt perceptual tasks (for illustration of these diametric models, see Fig. 1 of Gow et al., 2008; Fig 7: Myers and Blumstein, 2008). In attempts to resolve these conflicting models, Gow et al. (2008) used functional connectivity analyses applied to MEG data and showed that causal neural signaling directed from left SMG to “lower-level” areas (e.g., STG) modulates sensory representations for speech within a latency of 280–480 ms (Gow et al., 2008). The top-down nature of their effects strongly favored a feedback, perceptual account of the Ganong whereby lexical representations influence the earlier encoding of sublexical speech features (e.g., Myers and Blumstein, 2008; Noe and Fischer-Baum, 2020; van Linden et al., 2007).

Our EEG findings closely agree with MEG data by demonstrating a neural analog of Ganong biasing that unfolds very early in the chronometry of speech perception. We observed lexical modulation of speech ERPs beginning ∼200 ms after sound onset and no later than 300 ms. The early time window (200 ms) of these effects aligns roughly with the P2 wave of the auditory ERPs, a component that is highly sensitive to perceptual object formation, category structure (Bidelman et al., 2020; Bidelman et al., 2013; Bidelman and Walker, 2017; Liebenthal et al., 2010), and context effects in speech identification (Bidelman and Lee, 2015). Source analysis uncovered a Ganong neural circuit spanning five nodes including MTG, IPL, and MFG in left hemisphere and PCG, INS in right hemisphere. The engagement of frontal brain areas (MFG, INS) is consistent with the notion that lexical effects partly evoke post-perceptual, executive processes (Norris et al., 2000). The involvement of insular cortex is perhaps also expected in light of prior imaging work; bilateral inferior frontal activation is particularly evident for speech contrasts that are acoustically ambiguous (Bidelman and Dexter, 2015; Feng et al., 2018; Guediche et al., 2013) and under conditions of increased lexical uncertainty (Bidelman and Walker, 2019; Luthra et al., 2019) that place higher demands on attention (Bouton et al., 2018). Indeed, resolving phoneme ambiguity (as in the Ganong) may be one of the first processes to come online before the decoding of specific lexical features (Gwilliams et al., 2018). This may account for the early time course of our neural effects.

Notable among the Ganong circuit were nodes in left SMG and MTG. Critically, these regions were the only two areas associated with behavior illustrating their important role in the lexical effect. MTG forms a major component of the ventral speech-language pathway that performs sound-to-meaning inference and acts as a lexical interface linking phonological and semantic information (Hickok and Poeppel, 2007, 2004). MTG is also associated with accessing word meaning (Acheson and Hagoort, 2013), a likely operation in our Ganong task when ambiguous phonemes are perceptually (re)interpreted as words. Relatedly, left inferior parietal lobe an adjacent SMG are strongly recruited during auditory phoneme sound categorization (Desai et al., 2008; Gow et al., 2008; Luthra et al., in press), suggesting their role in phonological coding (Sliwinska et al., 2012). Parietal engagement is especially prominent when speech items are more perceptually confusable (Feng et al., 2018) or require added lexical readout as in Ganong paradigms (Oberfeld and Klöckner-Nowotny, 2016) and may serve as the sensory-motor interface for speech (Hickok et al., 2009; Hickok and Poeppel, 2000). Moreover, using machine learning to decode full brain EEG, we have recently shown that left SMG and related outputs from parietal cortex are among the most salient brain areas that code for category decisions (Al-Fahad et al., 2020; Mahmud et al., 2020). Similar results were obtained in a multivariate pattern decoding analysis of Luthra et al. (in press), who showed left parietal (SMG) and right temporal (MTG) regions were among the most informative for describing moment-to-moment variability in categorization. Additionally, the link between MTG and PCG implied in our data points to a pathway between the neural substrates that map sounds to meaning and sensorimotor regions that execute motor commands (Al-Fahad et al., 2020; Du et al., 2014). Still, the early time course of these neural effects (∼250 ms) occurs well before listeners’ behavioral RTs (cf. Fig. 2b vs. Fig. 4), suggesting these mechanisms operate at an early perceptual level. These findings lead us to infer that rapid (200-300 ms) context-dependent modulations within a restricted temporo-parietal circuit are most inducive to describing the degree to which listeners are susceptible to lexical influences during speech labeling.

Notably absent from our Ganong circuit—identified via differences waves—was canonical auditory-linguistic brain regions (e.g., STG). While somewhat unexpected, these data agree with previous fMRI results using a nearly identical Giss-Kiss continuum (Myers and Blumstein, 2008). Indeed, Myers and Blumstein (2008) reported strong IPL but no Ganong-related differences in several brain areas previously shown to be sensitive to phonetic category structure including STG and IFG. STG activity is greater when stimuli are maximally shifted from their VOT-matched counterparts (Myers and Blumstein, 2008). Although we observe a measurable Ganong effect, it is possible that stronger STG differentiation would have been observed in our EEG data with more salient lexical biasing stimuli. Still, the fact that correlations between neural and behavioral Ganong occurred in areas beyond canonical auditory-sensory cortex (e.g., STG) suggests that high order, top-down mechanisms drive or at least dominate lexical biasing (Gow et al., 2008) rather than auditory temporal cortex, *per se*. Though they do so rapidly. Alternatively, rather than a binary feedforward or feedback model of the lexical effect (Gow et al., 2008), it is possible the formation of speech categories operates in near parallel within lower-order (sensory) and higher-order (cognitive-control) brain structures (Mahmud et al., 2020; Toscano et al., 2018). Our data are broadly consistent with such notions. Category representations also need not be isomorphic across the brain. Category formation might reflect a cascade of events where speech units are reinforced and further discretized by a recontact of acoustic-phonetic with lexical representations (Mahmud et al., 2020; Myers and Blumstein, 2008).

Our data are best cast in terms of interactive rather than serial frameworks of speech perception as in the TRACE model of spoken word recognition (McClelland and Elman, 1986). As confirmed empirically (Ganong, 1980; Gow et al., 2008; Lam et al., 2017; Myers and Blumstein, 2008; Noe and Fischer-Baum, 2020), these models predict stronger lexical biasing when speech sounds carry ambiguity. Indeed, neural correlates of the Ganong effect were most evident at the midpoint of our speech continua, where word influences exert their strongest effect. The very nature of TRACE is that activation traverses from one level to the next before computations at any one stage are complete (McClelland and Elman, 1986). Indeed, available evidence coupled with present results suggest that word recognition could involve simultaneous activation of both continuous acoustic cues and phonological categories (Toscano et al., 2018). It is also possible that the acoustic–phonetic conversion and post-perceptual phonetic decision both localize so the same brain areas (Gow et al., 2008, p.621). Nevertheless, our data show that the acoustic-phonetic encoding of speech is rapidly subject to linguistic influences within several hundred milliseconds. While the early time-course implies a stage of perceptual processing, we find that lexical effects occur strongest outside the purview of canonical auditory-linguistic brain areas via a restricted temporoparietal circuit.

## Acknowledgments

We thank Dr. Emily Myers for sharing stimulus materials. This work was supported by the National Institute on Deafness and Other Communication Disorders of the National Institutes of Health under award number R01DC016267 (G.M.B.).

## Footnotes

1 EEG was not recorded from one participant due to technical error resulting in a final sample size of n=15 for the neural data (behavioral data were unaffected).

